# Conserved Genic Composition Unravels Rapid Karyotype Evolution and Polyploidization across Plants

**DOI:** 10.64898/2026.06.29.735181

**Authors:** Jianhui Gu, Wenli Chen, Dezhu Li, Haibao Tang, Xiang Li

## Abstract

Chromosomal variation underlies species evolution, but reconstructing its large-scale dynamics remains challenging, obscuring its adaptive significance. Here, we introduce *GouMang*, a framework that mines conserved genic section compositions across diverse species to trace karyotype evolution. In grasses, applied to 818 highly varied chromosomes spanning eight subfamilies, *GouMang* resolved a shared karyotype evolution path of 9 → 18 (ρ whole-genome duplication, ρWGD) → 12 chromosomes, followed by lineage-specific rearrangements or WGDs. Genes retained from the early ρWGD are linked to cold/light adaptation, supporting a key biomass-expansion event that impacted subsequent global ecological pattern and human agricultural civilization, in which K-Pg global cooling and subsequent forest degradation drove early understory grasses to sun plants. Parallel analysis in Brassicaceae reconstructed karyotype evolution as well as βWGD which unlinked to cold/light adaptation, reflecting a divergent biogeographic history compared to grasses. Together, *GouMang* depicts a widespread plant evolutionary pattern where karyotype constantly diversified with WGDs recurrently fueling adaptation.

## INTRODUCTION

Eukaryotic chromosomes function as the central platforms of genomic architecture, not only harboring protein-coding genes and regulatory elements ^1,2^ but also mediating DNA recombination and rearrangements with profound biological implications^3–6^. Whereas animal clades such as eutherian mammals typically maintain conserved karyotypes with moderate reshuffling^7,8^, angiosperms, a dominant component of terrestrial ecosystems, exhibit exceptional chromosomal dynamism, including rampant rearrangements and frequent WGD ^9–12^. Notably, chromosomal rearrangements have been linked to rapid adaptive radiation in several lineages, as demonstrated in cichlids from Lake Victoria ^13,14^. Since angiosperms even make Darwin so abominable by rapid radiation to dominate terrestrial niches (Darwin’s abominable mystery) ^15,16^, this raises the compelling question of how rampant karyotype variations contributed to the angiosperms and cereal evolutions.

Early efforts to clarify extensively-complex chromosome evolutions focused on grasses (Poaceae species) ^9–11^ due to that it contains numerous staple cereal crops and economically vital species that demanded genomic characterization. The initial “*top-down*” approach depended on aligning hundred low-density marker sequences among species, identifying syntenic chromosomal segments to deduce ancestral karyotypes ^9–11^. Results exhibits their frequent duplications and variable base chromosome numbers across grass lineages^17^, corresponding with its rapid evolution. Subsequent advancements in sequencing technologies enabled gene-resolution ancestral karyotyping in selected grass species ^12,18–21^. However, these studies were also limited by minor genome sampling, that missed most rearrangement features evolving over tens of millions of years. Further expanded sampling may still make ancestral proto-chromosome boundary ambiguities due to introduction of abundant chromosome rearrangements. Probably as such, the recent precise-gene-order approach AGORA ^22^ reconstructed high-resolution grass ancestral genome but not assessing karyotype *De Novo* (including chromosome variation).

A serial of another “*bottom-up*” approaches were also developed, that identifies exact rearrangement events between close-related species in multiple *bottom-to-up* cycles to deduces chromosome intermediates per each cycle ^23,24^. While “*bottom-up*” approaches excel at detecting individual rearrangement events, selecting species around evolutionary nodes could detect common rearrangements but remain challenging to tracing all complex rearrangements across lineage. With similar principle, an approach to construct phylogeny of lineages was developed by detecting evolutionarily conserved fusion breakpoints in a few of species (WGDI approach ^25–27^).

These current methods lack the capacity to trace complex karyotype evolution at high gene solution across large-scale species. To overcome this, we developed an approach *GouMang* (**<u>G</u>**enomic **<u>O</u>**rthology-**<u>U</u>**nified **<u>M</u>**ining of **<u>A</u>**ncestral **<u>N</u>**exus via **<u>G</u>**enome composition), by integrating precise gene-order reconstruction as well as a conserved chromosome components estimation. In the principle, since chromosomal rearrangement events may occur independently during evolution, species may simultaneously contain chromosomes identical to ancestral proto-chromosomes as well as those derived from inter-chromosome rearrangement events. Thus, the common chromosomes across large-scale species are more likely to represent ancestral proto-chromosomes, following the “Occam’s Razor” principle. *GouMang* contains six key analytical steps (Fig. 1a), including: (1) species-unified colinear block definition, (2) identification of evolutionarily conserved chromosomal compositions across wide species, (3) common chromosomes clustering, (4) ancestral genic karyotype reconstruction from both common and highly-rearranged chromosomal compositions, (5) subclade-specific karyotype dynamics evaluation, and (6) earlier pre-WGD genic karyotype determination - collectively overcoming the fundamental challenge of reconstructing complete karyotype evolutionary trajectories in rearrangement-prone taxa and thereby illuminating their adaptive roles during historical environmental upheavals.

**Fig. 1.**
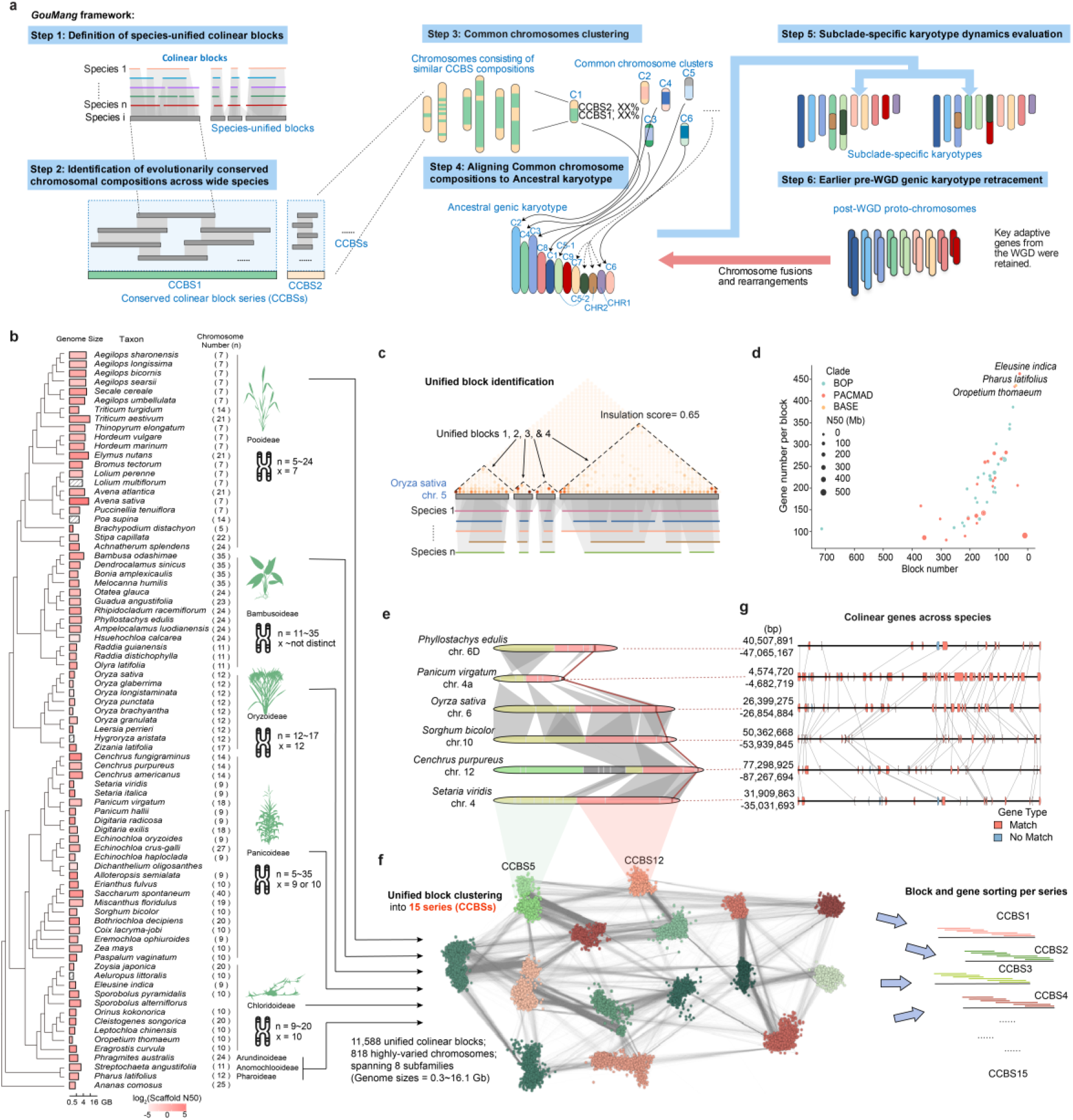
Unified block components of 818 high-varied chromosomes across grasses. **a**, The flow chart of *GouMang* framework including six analytical steps. **b**, Phylogenetic relationships of grass species included in this study, with genome sizes and chromosome numbers indicated for each. **c**, Example of unified block identification on a representative chromosome. **d**, Number and size distribution of unified blocks per species. **e**, Colinear overlaps of homologous chromosomal segments across species. Red lines indicate the positions of colinear genes shown in panel f. **f**, Clustering of all unified blocks based on synteny of inter-block overlaps. **g**, Detailed view of colinear gene pairs within a representative homologous segment.

## RESULTS

### Grass Genomes Include Extensive Chromosomal Variation

Poaceae represents a premier species reservoir in plant kingdom, comprising over 12,000 species with extreme karyotype diversity and dramatic genome size variation ^28–30^. We incorporated all 146 publicly available high-quality genome assemblies. Of them, 84 assemblies spanning 59 genera across eight main subfamilies (genome sizes = 0.3-16.1 Gb) are near-chromosome-scale, enabling precise detection of chromosomal rearrangements (Fig. 1b; Supplementary Table 1). Two species in the basal subfamilies Anomochlooideae and Pharoideae are collected with n=11 and n=12 chromosomes, respectively. Among the sampled modern subfamilies, Pooideae demonstrates a large chromosome number variation (n=5-24 across 22 species), while most species retain the x=7 karyotypes; Bambusoideae shows remarkable chromosomal plasticity (n=11-35 across 13 species) without a predominant base number; Oryzoideae maintains relative stability (n=12-17 across 9 species) with x=12 as the most common base number; Panicoideae exhibits substantial diversity (n=7-40 across 23 species), with most tribes showing a clear x=9 or x=10 predominance; Chloridoideae (n=9-20 across 10 species) predominantly maintain x=10 chromosomes. In the smaller subfamily Arundinoideae, *Phragmites australis* (reed) genome was collected. Though the sequencing individual exhibited n=24, individuals with varying ploidies have been documented in surveys^31^. This set represents a microcosm of Poaceae diversity, with balanced sampling to evolutional taxon and sufficient chromosomal variation.

### Pan-Grass Unified Colinear Block Definition

The initial step of *GouMang* pipeline defines unified colinear blocks for each species (Fig. 1c), addressing a critical limitation that generate inconsistent blocks when aligning the same genome to different reference species due to their reliance on variable orthologous gene anchors. Pairwise collinear blocks across all possible species combinations were identified, then unified block definitions were established by selecting the most frequently observed boundaries supported by high insulation scores - a well-established metric also used for chromatin domain demarcation ^32^. To avoid overrepresented selection from the same genus and thus bias ancestral karyotype, we implemented a subsampling strategy (four species per genus, thus 54 species) for unified block clustering while retaining all species for subsequent ancestral karyotype assessment and rearrangement analysis. This simple design avoids repeated reclustering when new genomes are added, especially for the rapidly expanding genomic datasets (See Discussion). In these samples, 1,157,105 initial pairwise collinear blocks were identified spanning 16,712 orthologs, with maximum 1,902 orthologs per block. Removing microsyntenic units (≤10 genes) to reduce noise from stochastic rearrangements or paralogous genes, 484,050 high-confidence collinear blocks were retained. Along each chromosome, we computed insulation score metric of collinear blocks across all pairwise-species comparisons. A high insulation score (≥0.65) points to distinct boundaries along chromosome, median 79.79% cross-species block definitions, delineating 11,588 unified colinear blocks from 818 chromosomes. For example, rice chromosome 5 could be partitioned into four unified colinear blocks, supported by over 78.87 % of collinear blocks from other species (Fig. 1c).

The number of unified colinear blocks per species ranged from 28 to 713 (median: 137), with 75.9% of species containing <200 blocks (Fig. 1d). Block consisted 78 orthologs in median, with maximum 1,622 orthologs. A striking pattern emerged: species with >200 blocks consistently showed <200 genes per block and possessed larger genomes. This trend peaked in *Triticum aestivum* with 713 blocks (the maximum observed), merely 106 genes per block (minimum density), and a 14.2 Gb genome. This suggests that genome expansion correlates with increased block rearrangements. Conversely, species with smaller genomes like *Eleusine indica* (28 blocks, 462.75 genes per block) retained fewer blocks with more genes per block, likely representing closer architecture of preserving ancestral segments with less rearrangements.

### Pan-Grass Unified Colinear Blocks Exhibit 15 Conserved Series

The unified blocks distributed along homologous chromosomes, such as those in rice chr. 6, *Phyllostachys edulis* chr. 6D, *Panicum virgatum* chr. 4a, *Sorghum bicolor* chr. 10, *Cenchrus purpureus* chr. 12, and *Setaria viridis* chr. 4, exhibit extensive syntenic overlap (Fig. 1e). This observation led us to hypothesize that frequent adjacent segments in most genomes could be identified based on synteny values of inter-block overlaps. We therefore performed unsupervised clustering using the synteny Jaccard index (≥ 0.7), which grouped all unified colinear blocks into 15 conserved collinear block series (CCBSs) (Fig. 1f). In these representative chromosomes (Fig. 1e), the overlapping blocks are encompassed by two distinct CCBSs (CCBS5 and CCBS12), despite *Cenchrus purpureus* chr. 12 contains a third CCBS composition fused to these two. These results collectively demonstrate that the highly diversified grass genomes are composed of a limited number of similar chromosomal compositions.

Genes were reliably anchored within fine-grained segments, as illustrated by the small syntenic blocks in representative chromosomes (Fig. 1g; also indicated in red colinear lines of Fig. 1e). This fine-scale resolution enabled the reconstruction of consensus gene orders within CCBSs. Using a length-priority block ordering strategy analogous to AGORA ^22^, we inferred consensus gene joins from the most frequent adjacencies across species (Fig. 1f). Each CCBS comprises 3,336-7,527 gene joins (median: 3,806; mean: 4,049), with individual joins supported by median 85% of species. Crucially, join support frequencies showed no correlation with genome assembly N50 (Spearman’s *ρ* = 0.116, *P* = 0.31), confirming that the observed diversity of gene rearrangements reflects true evolutionary divergence rather than false positives arising from assembly artifacts.

### Nine Common Chromosome Groups Across Grass Genomes

Then, we reconstructed ancestral karyotype based on an evolutionary principle: given that chromosomes rearranged independently across lineages, species contain chromosomes with ancestral segment compositions and new inter-chromosomal rearrangement events. Consequently, across lineages, chromosomes retaining ancestral segment compositions appear with higher frequency than any specific rearrangement variants (Fig. 2a). We therefore proposed most chromosomes exhibiting universally conserved CCBS patterns shared among multiple lineages as candidate ancestral components.

**Fig. 2.**
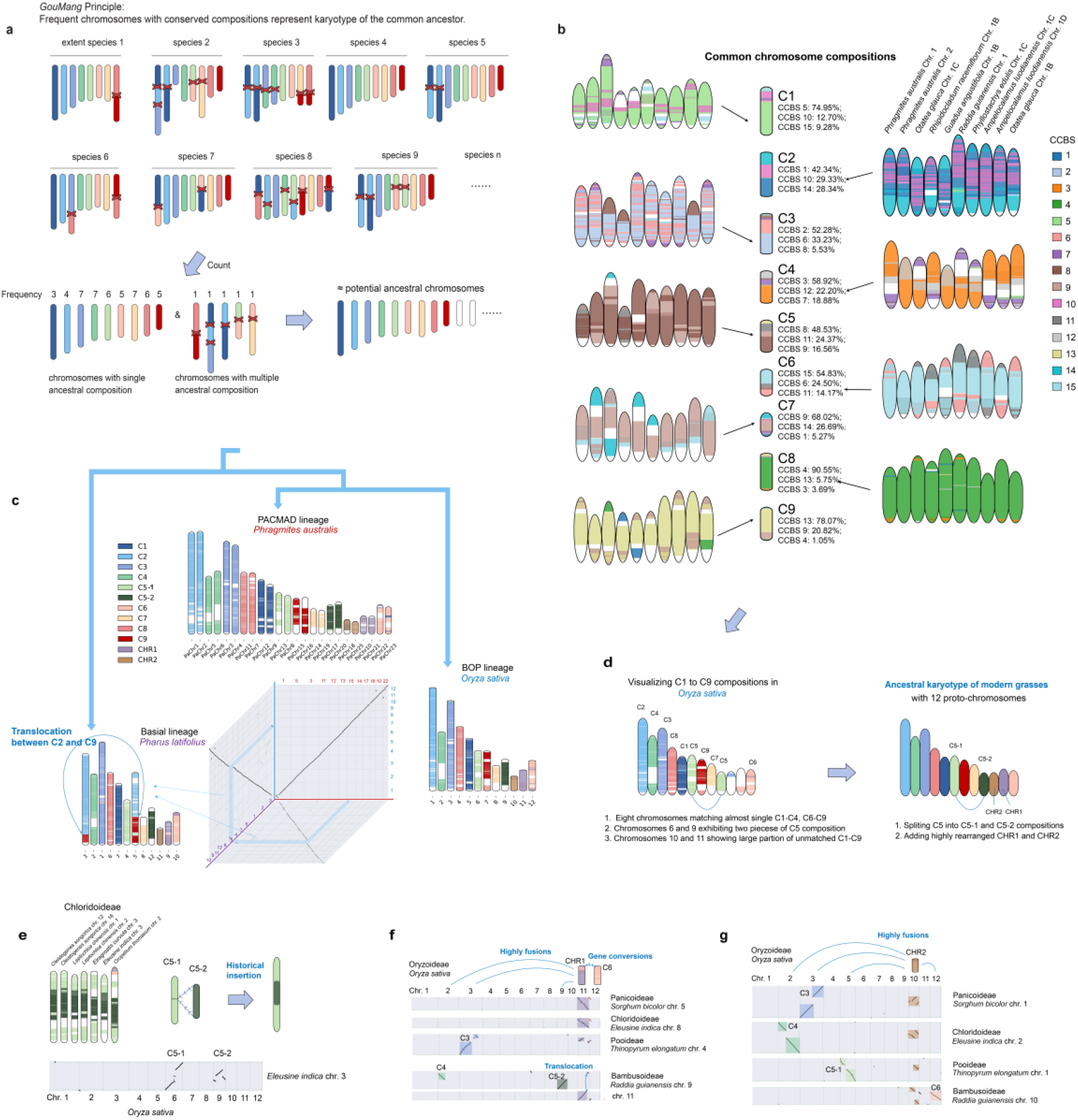
Ancestral karyotype reconstruction reveals conserved and rearranged chromosomal components. **a**, *GouMang* principle Schematic for identifying evolutionarily conserved chromosomes across large-scale speices. **b**, Block compositions of the nine common grass chromosome groups, clustered by using CCBS≤2 orthologs. **c**, Similar karyotypes and genome synteny among distinctly-related species from early-diverging grass lineages, PACMAD, and POB clades. **d**, Reconstruction of the ancestral 12-chromosome karyotype in modern grasses. **e**, Shared insertions of C5-2 into C5-1 in Chloridoid. **f**, Evolutionary rearrangements and gene conversion events involving CHR1 across major grass subfamilies. **g**, Evolutionary rearrangements involving CHR2 across major grass subfamilies.

Given the high frequency of chromosomal duplications and rearrangements during Poaceae evolution, clustering chromosomes by their CCBS composition still remains not simple, probably making ambiguous clusters. This is corresponding to an observation that the 15 CCBSs contains totally 60,735 consensus gene joins, excess over threefold higher than the average ortholog content per species (~16,712), implying that a large number of orthologs anchored into distinct CCBSs. Therefore, removing orthologs anchored at multiple CCBSs may clarify the chromosome clusters. To ensure higher gene resolutions in subsequent ancestral karyotype, we defined ranging sets of orthologs anchored across number-various CCBSs (from one to multiple) (Extended Data Fig. 1a) and estimated which set represents low inter-cluster gene similarity (Extended Data Fig. 1b) and high genome coverage (Extended Data Fig. 1c).

In results, chromosome clusters using orthologs in no more than two CCBSs provides the most reliable reconstruction of Poaceae conserved chromosomes (Fig. 2b), based on the three evidences: 1) Using exclusively single-CCBS orthologs, inter-cluster gene similarity was minimized (1%), while intra-cluster similarity peaked (89%). Gradually adding multi-CCBS orthologs increased inter-cluster similarity (maximum, 10% at orthologs anchored in no more than 7 CCBSs) and declined intra-cluster similarity (minimum, 35% at orthologs anchored in no more than 7 CCBSs), demonstrating ambiguous chromosomes clustering (Extended Data Fig. 1b). However, 2) selecting only single-CCBS orthologs exhibits an excessively low genome coverage (median 27%, maximum 64%), lacking a representation for whole chromosomal segments (Extended Data Fig. 1c). Adding two-CCBS orthologs has shown sufficient genome coverages for assessing whole chromosome variation (median 65%, maximum 91%), while intra- (72%) and inter- (3%) cluster similarity of this set remains closed to the single-CCBS orthologs’ set, serving as best selection to come over the ambiguity-coverage compromise. 3) CCBS compositions for this set clustered 682 chromosomes as nine groups (C1 to C9 components), exhibiting distinct CCBS compositional patterns across clusters but striking conservation within each cluster (Fig. 2b). We also test the chromosome set by using CCBS≤3 orthologs, also resulting in nine clusters but more similar composition between clusters (specially for C4 & C5) (Extended Data Fig. 2), suggesting that using CCBS≤2 orthologs may clarify chromosome cluster while adding more-CCBS orthologs make clusters ambiguous. These results confirm not only nine common chromosome groups with similar CCBS compositions per chromosome (Fig. 2b) but also extensive rearrangements in Poaceae.

### A 12-Chromosome Genic Karyotype in Common Ancestral of Extant Grasses

Karyotyping all chromosomes of the extant species by using orthologs of the nine common chromosomes (Extended Data Figs. 3-7), we observed a high frequency of recombination between C1 to C9 components. Despite this, a conserved karyotype shared among BOP species like rice and PACMAD species like reed (one subgenome) were discovered, consisting of 12 chromosomes (Fig. 2c), accompanying with a high synteny between homologous chromosomes of the two species. Comparing to a karyotype of the basial lineages’ representative, *Pharus latifolius*, which also contains 12 chromosomes, the rice and reed karyotypes display only one translocation event between C2 and C9 components (Fig. 2c), proposing a 12-chromosome karyotype that likely originated in a common ancestor of modern grasses.

In rice karyotype by C1 to C9 compositions, eight chromosomes maintaining full C1, C2, C3, C4, C6, C7, C8, and C9 compositions, two chromosomes (chrs. 6 and 9) corresponding to segmented C5, and two chromosomes (chrs. 10 and 11) almost lacking C1-C9 homology (Fig. 2d). To explain why not all chromosomes align to full common chromosome components, these mis-aligned chromosomes were compared side-by-side across lineages: 1) For C5 segments (Fig. 2e), components aligning chrs. 6 and 9 in rice were further divided into C5-1 and C5-2, respectively. Re-aligning them across grasses, we observed that C5-2 components in Chloridoideae fused C5-1 segments at two ends, suggesting an event of C5-2 chromosome insertion (Fig. 2e), while the two components in other subfamilies exhibit unconnected. This may cause the mix of C5-1 and C5-2 components in primary clustering. 2) For chr. 11 in rice (Fig. 2f), a minimal partial was aligned to C6 (one end of chr. 12), which was previously described as a species-shared but level-varying gene conversion between early-duplicated chromosomes, analogous to those reported between rice chrs. 12 and 11, and sorghum chrs. 8 and 5 ^33,34^. Besides, in Pooideae, *Thinopyrum elongatum* retains only 70% of the syntenic segments compared to rice chr. 11, indicating a 30% compositional loss, and has fused with the C3 composition to form chr. 4. Similarly, in Bambusoideae, the homologous region of rice chr. 11 underwent translocation, with a small segment relocated to a fused chromosome consisting of C4 and C5-2. These observations consistently highlight the highly rearranged nature of this chromosomal component, which we designate as highly rearranged component 1 (CHR1). 3) The homologous segments of rice chr. 10 exhibit extensive fusion events with different chromosomal components across major grass subfamilies (Fig. 2g): in *Sorghum bicolor* (Panicoideae) it is fused with C3, in *Eleusine indica* (Chloridoideae) with C4, in *Thinopyrum elongatum* (Pooideae) with C5-1, and in *Raddia guianensis* (Bambusoideae) with C6. These consistent yet divergent fusion patterns underscore the highly rearranged nature of this component, which we designate as highly rearranged component 2 (CHR2). Based on these findings, we propose that the ancestral karyotype of modern Poaceae likely comprised twelve proto-chromosomes, including C1, C2, C3, C4, C5-1, C5-2, C6, C7, C8, C9, CHR1 and CHR2. (Fig. 2d).

We then detected the retainment of individual (not fused) ancestral components in species, to test the *GouMang* principle that common chromosomes represent most ancestral ones. While not every species retains all twelve unfused ancestral chromosomal components, a side-by-side comparison across species reveals complete set of unfused ancestral chromosomal components, albeit with a subfamily-preferred distribution pattern (Fig. 3a). For instance, Oryzoideae and Bambusoideae (e.g. Phyllostachys edulis) with higher chromosome base number (dominant x = ~12) retain more unfused ancestral chromosomal components, while Pooideae (dominant x = ~7) contained only three preferred unfused common components (C2, C3, and C4) (Fig. 3a). This may suggest that chromosome number constraint was associated with ancestral chromosomal fusions. An extreme case is *Brachypodium distachyon* that has 5 chromosomes, where all ancestral chromosomes except C8 (chr. 5) were fused. Another species, *Zea mays*, undergoing WGD but retaining a pre-WGD chromosome number (n=10) ^9,11^, also show lacking unfused all ancestral components, while its closely related species *Sorghum bicolor* (n=10) contains eight unfused ancestral components.

**Fig. 3.**
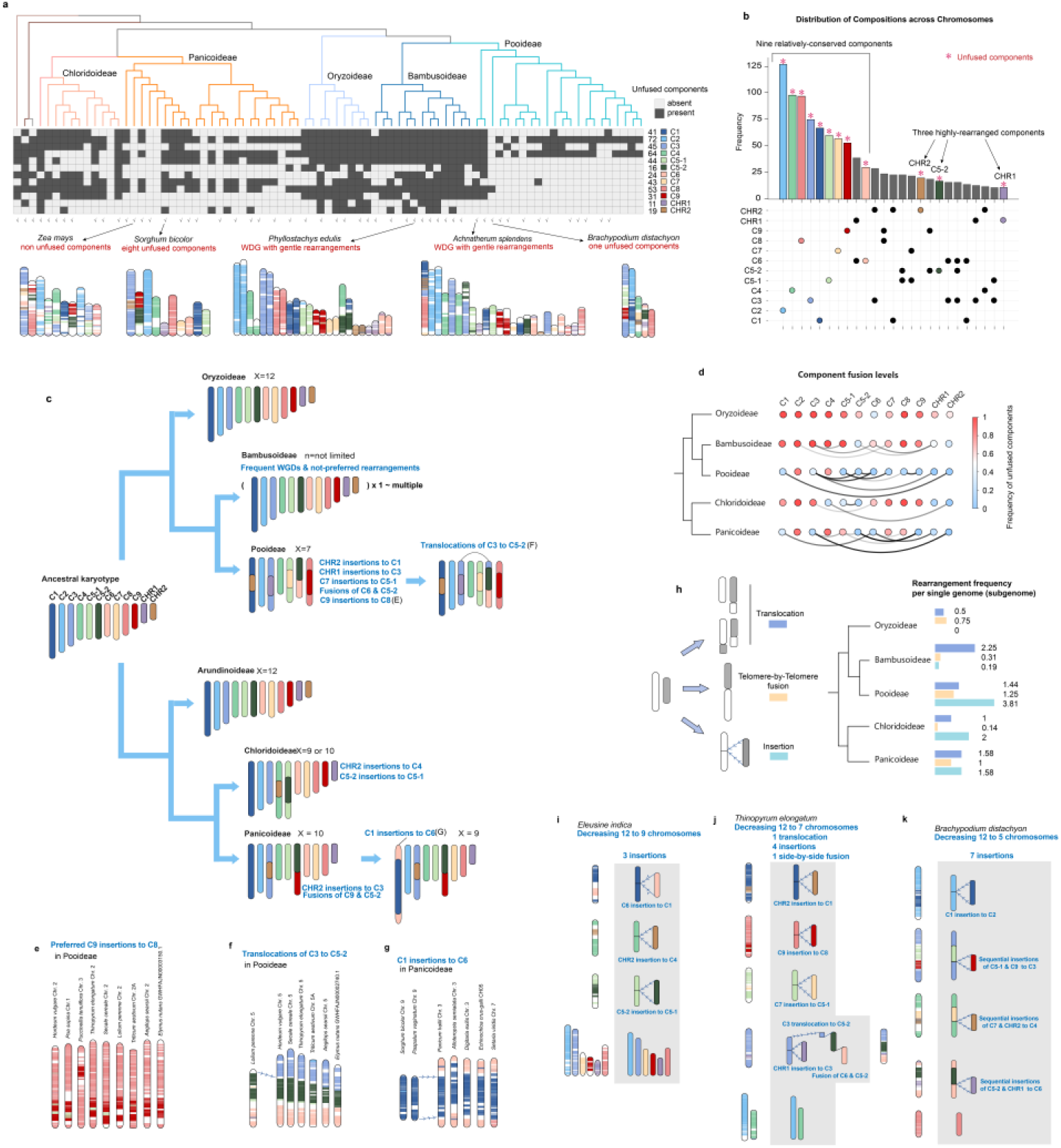
Divergence in chromosomal evolution across grass subfamilies. **a**, Distribution of unfused chromosomal components across species of each subfamily. Numbers on the right indicate the count of species containing the corresponding component. Checkmarks at the bottom denote species used for *de novo* reconstruction of the 12-chromosome ancestral karyotype; unmarked species were karyotyped directly. The two sets exhibit consistent patterns of unfused components within subfamilies, supporting the reliability of our approach. **b**, Counts of chromosome with single or rearranged components across all species, revealing nine relatively conserved and three highly rearranged components. **c**, Schematic representation of karyotype diversification, illustrating subfamily-preferred karyotypic configurations. **d**, Fusion frequency among components across subfamilies. Thicker connecting lines indicate higher fusion frequency. **e**, Preferred insertions of C9 into C8, shared among many species in the Pooideae. **f**, Preferred translocations between C3 and C5-2 in Pooideae. **g**, Preferred insertions of C1 into C6 frequently observed in the Panicoideae. **h**, Frequency of different rearrangement types in each subfamily. **i-k**, Examples of three species demonstrating how rearrangements led to chromosome reduction from the ancestral 12-chromosome karyotype.

An analysis of chromosome composition across grasses (Fig. 3b) revealed that unfused C2, C4, C8, C3, C1, C5-1, C7, and C9 components were the eight most frequently retained elements. Fused chromosomes consisting of C6 and CHR1 ranked ninth, followed by unfused C6 chromosomes in tenth place. The high frequency of C6-CHR1 fusion may be attributable to species-shared but quantitatively variable gene conversion events (Fig. 2f). Except these, unfused CHR1, C5-2, and CHR2 components ranked 16th, 18th, and 25th, respectively. Together, these results support an ancestral karyotype consisting of nine evolutionarily conserved and three highly rearranged components, and confirm the reliability of our reconstruction through identifying common chromosomal compositions.

### Karyotype Divergence towards Extant Grass Lineages

We next deeply analyzed subfamily-preferred rearrangements (Fig. 3c), through comparing karyotypes of all species (Extended Data Figs. 3-7) and estimating the fusion levels of 12 ancestral chromosomal components in each subfamily taxa (Fig. 3d). In the BOP clade, Oryzoideae exhibits less C1-C9 fusions with ancestral-like chromosome number (n = 12) (Extended Data Fig. 3). Bambusoidea exhibits frequent WGDs but a limited level of fusions for species (Extended Data Fig. 4), like between C3, C5-1, C6, and CHR1 components (Fig. 3d), supporting also low derivation from the 12-chromosome ancestral karyotype. In Pooideae (Extended Data Fig. 5), despite the basial lineages like *Achnatherum splendens* (n = 24) contain nine individual ancestral chromosomal components (Fig. 3a), most species constrain chromosome number per subgenome as 7 (only 2-3 individual ancestral chromosomes) (Fig. 3c), with particularly extensive fusions, including CHR2 insertions to C1, CHR1 insertions to C3, C7 insertions to C5-1, C9 insertions to C8, telomere-by-telomere fusions of C6 and C5-2, and subsequent translocations of C3 to the C5-2 & C8 fused chromosomes. The chromosome patterns involving Pooideae-specific “C9 insertions into C8” (Fig. 3e) and “C3 translocations” (Fig. 3f) were specifically highlighted to illustrate karyotype divergence. The fusion levels of ancestral chromosomal components in Pooideae also support their karyotype divergence (Fig. 3d).

In the PACMAD clade, Chloridoideae (Extended Data Fig. 6) exhibits chromosome number x=9 or 10 per genome, with preferred CHR2 insertions to C4 and C5-2 insertions to C5-1 (above described in Fig. 2e). Panicoideae (Extended Data Fig. 7) exhibits a high frequency of CHR2 insertions into C3, telomere-by-telomere fusions of C9 and C5 components, and partial-species C1 insertions to C6. The chromosome patterns involving the C1 insertions into C6 were plot to highlight karyotype dynamism (Fig. 3g). Beyond these, abundant rearrangements were also validated by the detections of fusion levels among ancestral chromosomal components (Fig. 3d), showing diverse chromosome variations across lineages.

### Subfamily-Preferred Insertions and Fusions Decrease Chromosomes

To further characterize the rearrangement preference underlying karyotype divergence, we estimated the rearrangement types in each subfamily, including chromosomal translations (composition recombinations without chromosome number changing), telomere-by-telomere fusions (composition recombinations with chromosome reduce), and insertions (one composition surrounding another composition with chromosome reduce) (Fig. 3d). The Oryzoideae subfamily exhibits relatively low levels of translocations (0.5 per genome/subgenome) and telomere-by-telomere fusions (0.75), suggesting a slower karyotype evolutionary rate, which may explain its conserved x = 12 chromosome number and close resemblance to the ancestral karyotype. In contrast, Bambusoideae shows a high translocation frequency (2.25) but very low rates of telomere-by-telomere fusions (0.31) and insertions (0.19), consistent with its history of genome duplication without substantial chromosome number reduction. Pooideae displays near-average translocation rates (1.44) but the highest levels of telomere-by-telomere fusions (1.25) and insertions (3.81) among all subfamilies, correlating with its extensively reduced chromosome numbers (dominant x = 7). Within the PACMAD clade, both Chloridoideae and Panicoideae show moderate translocation frequencies (1.0 and 1.5, respectively), with sum of fusion and insertion frequencies ranging between 2 and 3 events per genome, consistent with their characteristic base chromosome numbers of 9 to 10.

Furthermore, we examined three representative lineages with markedly reduced chromosome numbers, including *Eleusine indica* (Chloridoideae; n = 9), *Thinopyrum elongatum* (Pooideae; n = 7), and *Brachypodium distachyon* (Pooideae; n=5), to validate the role of rearrangements in chromosome number reduction. *Eleusine indica* (Fig. 3i) underwent three insertion events, incorporating C6, CHR2, and C5-2 into C1, C4, and C5-1, respectively, reducing its chromosome number by three (from 12 to 9). Similarly, *Thinopyrum elongatum* (Fig. 3j) experienced four insertion events and one telomere-by-telomere fusion, collectively reducing the chromosome count by five (to n=7), along with one translocation that did not alter chromosome number. In contrast, *Brachypodium distachyon* (Fig. 3k) underwent a distinct set of probably up to seven insertions, including one C1 insertion into C2 (forming its chromosome 2) and three sequential-insertion chromosomes, where C5-1 and C9 components inserted into C3, C7 and CHR2 into C4, as well as C5-2 and CHR1 into C6. This is similar to the previously-reported sequential insertions ^35^.

These karyotypic analyses confirm that chromosomal number reduction in grasses occurred primarily through insertions and telomere-by-telomere fusions, with insertions being the dominant mechanism. Notably, no evidence was found for chromosomal splits, suggesting that increases in chromosome number in grasses occur predominantly through whole-genome duplication rather than fission events. Based on these findings, we proposed that karyotype evolution is driven by cyclical WGDs followed by lineage-specific rearrangements, with insertions and fusions serving as the key mechanisms for chromosomal reduction.

### Nine Earlier Proto-Chromosomes of Pre-ρWGD Karyotype

To trace the origin of the ancestral 12-chromosome karyotype, we analyzed its derivation from the pre-ρWGD karyotype through comparative synteny within single genomes. The C1-C9-CHR compositions consistently aligned into five syntenic groups across species (Figs. 4a and 4b). For instance, in rice (Oryzoideae, n=12), (Group 1) chromosomes 1 (belongs to C2) and 5 (C1) showed a pairwise synteny; (Group 2) chromosome 3 (C3) are colinear to combinations of whole chromosomes 7 (C9), 10 (CHR2), and partial chromosome 12 (C6); (Group 3) chromosome 2 (C4) are colinear to combinations of partial chromosome 4 (C8) and whole chromosome 6 (C5); (Group 4) chromosome 8 (C7) is colinear to combinations of partial chromosome 4 (C8) and whole chromosome 9 (C5); (Group 5) chromosomes 11 (CHR1) is colinear to partial chromosome 12 (C6) (Fig. 4a). Similar patterns were observed in sorghum (Panicoideae, n=10), confirming the conservation of these syntenic groups despite differences in chromosome number (Fig. 4b).

**Fig. 4.**
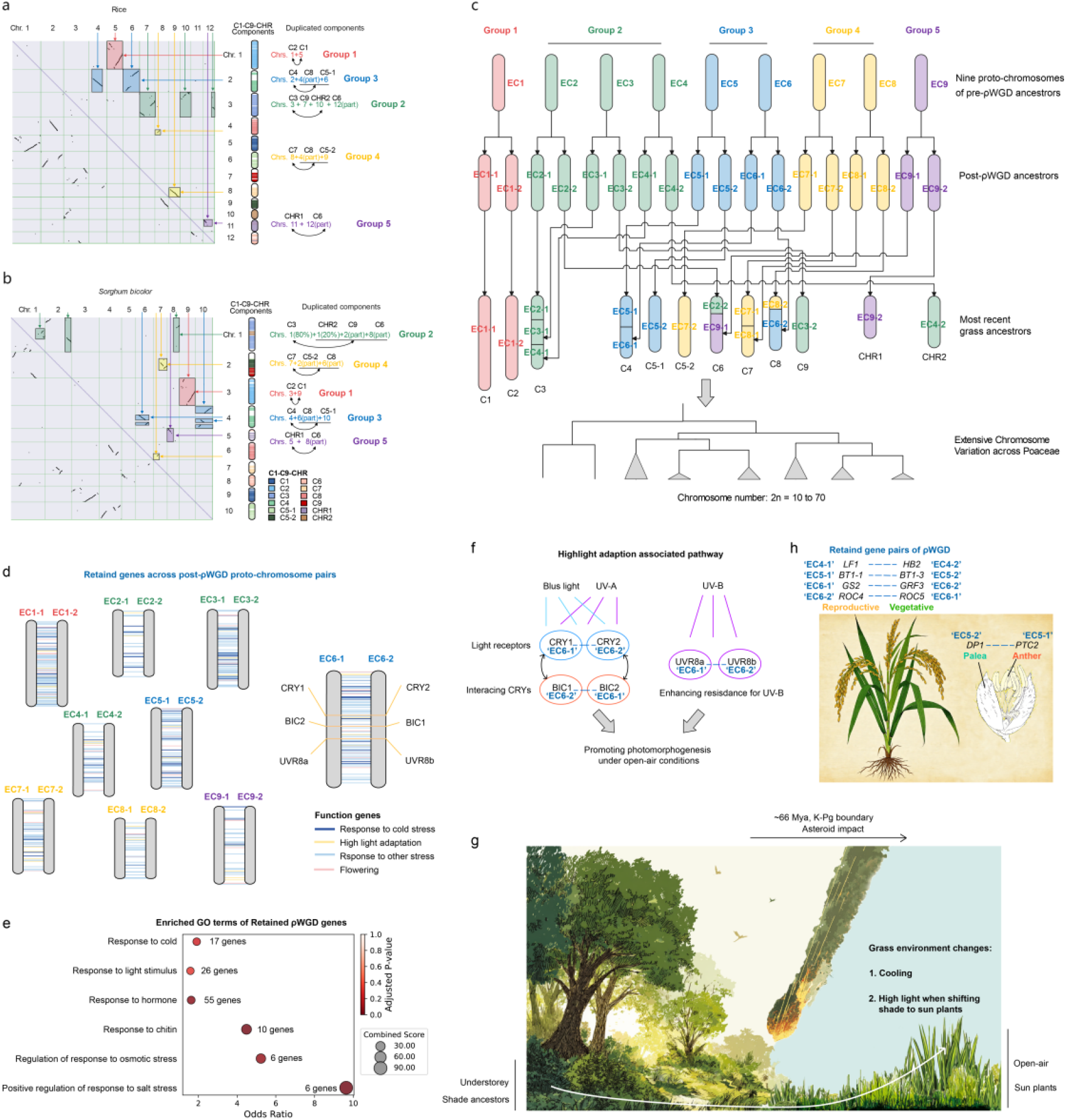
Karyotype dynamics and functional legacy of ρWGD in grass evolution. **a-b**, Intra-genomic synteny analysis in rice (a) and sorghum (b), identifying five conserved colinear groups. **c**, Karyotype changes before and after the ρWGD event in the ancestral grass lineage. **d**, Retained genes across the nine proto-chromosomal WGD duplicate pairs. **e**, Representative GO enrichments of retained genes associated with stress responses. **f**, Genes implicated in high-light adaptation retained after ρWGD. **g**, Proposed scenario of early grass habitat transition and environmental adaptation. **h**, Retained duplicate gene pairs showing functional divergence in vegetative growth and plant architecture regulation.

The pervasive intra-genome synteny between duplicated chromosomal segments confirms a Poaceae-shared ρWGD, an event that may help adapt to the climatic transitions during K-Pg boundary (~66 Mya) ^30^. We estimated the maximum chromosome number of pre-ρ WGD karyotype as nine (EC1-EC9) and traced the fusion history of post WGD chromosomes to form 12 common chromosomes of Poaceae ancestors (Fig. 4c). Based on pairwise synteny of C1 and C2, we traced them from a duplication (EC1-1 and EC1-2) of one earlier chromosome (EC1) without inter-chromosomal fusions. The group 2 may incorporate three earlier chromosomes (EC2, EC3, and EC4), but did not serve as one proto-chromosome. If only one earlier chromosome existed, one post-WGD chromosome would evolve to C3, while another post-WGD chromosome split to the CHR2, C9, and partial C6 (Extended Data Fig. 8). This conflicts with the evolutionary features not preferring chromosomal splits (See above). Instead, we proposed that three individual chromosomes EC2, EC3, and EC4 may duplicate, then single copies of the post-WGD chromosomes (EC2-1, EC3-1, and EC4-1) fused to C3, while the other copies evolved to CHR2 (EC2-2), C9 (EC4-2), and partial C6 (EC3-2) (Fig. 4c). Similarly, the group 3 may include two earlier chromosomes (EC5 and EC6). Single copies of the post-WGD chromosomes (EC5-1 and EC6-1) fused to C4, while the other copies evolved to C5-1 (EC5-2) and partial C8 (EC6-2). The group 4 may also include two earlier chromosomes (EC7 and EC8). Single copies of the post-WGD chromosomes (EC7-1 and EC8-1) fused to C7, while the other copies evolved to C5-2 (EC7-2) and partial C8 (EC8-2). The group 5 contains only one earlier chromosome (EC9), comprising the post-WGD chromosomes in the last chromosomal segments partial C6 (EC9-1) and whole CHR1 (EC9-2) (Fig. 4c). These results revealed the evolutionary trajectory that a nine-chromosome earlier karyotype duplicated and rearranged to form a 12-chromosome karyotype of most recent Poaceae ancestors.

Moreover, the karyotype of the basial lineages’ representative, *Pharus latifolius*, shows limited variance (only one translocation between C2 and C9) from the reconstructed 12-chromosome ancestral karyotype of modern grasses (Fig. 2c), supporting the view that the all Poaceae lineages, including the basal groups, show a common ancestor of 12-chromosome ancestral karyotype (C1-C9-CHR), which were derived from the ρWGD and subsequent fusions of nine-chromosome karyotype (EC1-EC9).

### ρWGD Drove Grass to Sun Plant, Key for Crop and Herbivore Ecosystem Rise

Given broad consensus that WGDs around K-Pg boundary have potentially helped species survive the extinction event ^36,37^, we analyzed retained ρWGD genes to evaluate their contribution to the origin and adaptive evolution of grasses. Although many gene copies are lost after WGD due to relaxed selection, some are retained and can contribute to functional innovation or pathway complexity ^38,39^. We identified 3,700 retained duplicated gene pairs distributed across nine duplicated chromosomal components (Fig. 4d; Supplementary Table 2), with serials of enriched Gene Ontologies (GOs) involved in fundamental pathways (Supplementary Table 3). The top significant GO terms represent fundamental regulatory processes, including DNA-binding transcription factor activity (GO:0003700), transcription regulator activity (GO:0140110), regulation of DNA-templated transcription (GO:0006355), regulation of RNA biosynthetic process (GO:2001141), and regulation of RNA metabolic process (GO:0051252). These findings are consistent with previous reports indicating that dosage-sensitive genes, particularly those whose imbalance may disrupt regulatory networks, are often selectively retained ^38,39^. This confirmed the reliability of our retained gene analysis.

Beyond these, multiple stress-response pathways were also significantly enriched (Fig. 4e; Supplementary Table 3), including response to cold (GO:0009409, 17genes), light stimulus (GO:0009416, 26 genes), hormone (GO:0009725, 55 genes), chitin (GO:0010200, 10 genes), salt (GO:1901002, GO:1901000, 6 genes), and osmotic stress (GO:0047484, GO:0006972, 6 genes). This enrichment supports the view that grasses experienced collective severe environmental transitions following the ρWGD, consistent with the complex changes documented during the K-Pg boundary, including global cooling, darkness, acid rain, and widespread wildfires ^40^. The pronounced retention of cold-response genes were parallelly found in multiple angiosperm lineages that underwent contemporaneous WGDs ^41^, indicating that cooling constituted a major selective pressure on early grasses and other angiosperms.

In contrast to previous reports that WGDs facilitated angiosperm lineages to adapt to the low-light conditions caused by global dust clouds after the asteroid impact during the K-Pg boundary ^40,41^, the ρWGD-retained gene pairs which we found in grasses do not contain the darkness-related genes such as phytochromes (e.g., *PHY*) as the multiple angiosperm WGDs retained. Meanwhile, we re-examined that grass ρWGD-retained genes in the published datasets^41^ exclude darkness-related *PHY* genes, which present in the other angiosperm lineages. Furthermore, our dataset uncovered a significant enrichment of response pathway to light stimulus (GO:0009416, 26 genes) and identifications of several retained genes associated with high light adapted photomorphogenesis at the proto-chromosome pair EC6-1 & EC6-2 (Fig. 4d), including: cryptochromes *CRY1* and *CRY2*, which sense blue/UV-A light and promote photomorphogenesis ^42^; their repressors *BIC1* and *BIC2* that fine-tune CRY activity ^43^; and UV-B receptors *UVR8a* and *UVR8b*, which mitigate UV-B inhibition of growth ^44,45^ (Fig. 4f). These results suggest that early grasses likely encountered high-light stress rather than darkness during the K-Pg transition.

Integrating the phylogenetic evidence ^46–48^ that early-diverging grass lineages (such as *Pharus latifolius*) occupied shaded understory habitats and most modern grasses (e.g., rice and maize) thrive in open, high-light environments, we proposed a revised narrative (Fig. 4g): while many sun-adapted plants faced severe low-light stress due to impact-induced dust clouds, early grasses, thus already adapted to understory shaded conditions, were likely less affected by reduced light levels. Instead, global cooling and consequent forest degradation may have forced these understory grasses into open habitats, driving their adaptation to cold and higher light intensities. Our results do not challenge existing models of WGD-enabled adaptation to darkness and cooling in most plants, but rather illuminate a distinct evolutionary trajectory in grasses, an original lineage living in understory.

Considering that shade-adapted plants could not have achieved the high biomass/yield from the capture of insufficient solar energy, the transition to sun plants likely represents the key trigger for the emergence of dozens of cereal crops within grasses ^49^ and the massive evolutionary radiation of herbivore lineages ^50^. We proposed that the ρWGD retained genes involved in light stimulus and photomorphogenesis constitutes the key genetic basis for this transformative shift.

Furthermore, the retained gene pairs also include several those undergoing functional divergence between reproductive and vegetative development. For instance, key genes such as *LF1, BT1-1, GS2*, and *ROC4* play more critical roles in reproductive growth and crop yield, while their respective paired duplicates, including *HB2, BT1-3, GRF3*, and *ROC5*, are predominantly involved in vegetative growth and plant architecture regulation (Fig. 4h). We also identified the *DP1* and *PTC2* gene pair, in which *DP1* primarily functions in the palea, while *PTC2* is mainly expressed in the anther. The functional divergence of these retained gene pairs likely contributed crucially to the subsequent formation and domestication of modern crops.

### Dramatic Karyotype Evolution as well as βWGDs in Brassicaceae

Parallelly, we assessed karyotype evolution diversity across angiosperm lineages through integrating 38 genomes of a large dicot family Brassicaceae with highly varied chromosome numbers (n = 5-20) spanning six subfamilies (Fig. 5a; Supplementary Table 4). Ten CCBSs covering 92.17% of unified blocks were grouped into five common chromosomal components (C1-C5) (Fig. 5b). Aligning these components to all chromosomes revealed widespread inter-component rearrangements (Extended Data Fig. 10). Strikingly, *Capsella orientalis* and *Arabidopsis lyrata* (Camelinodae) retained eight chromosomes without inter-component rearrangements and component duplications (Fig. 5c). We next disentangled these C2 and C3 into distinct components residing on separate chromosomes of *Capsella orientalis*, designating them C2-1/C2-2 and C3-1/C3-2/C3-3 (Fig. 5c). Side-by-side analysis shows that the eight chromosome groups within single components (C1, C2-1, C2-2, C3-1, C3-2, C3-3, C4, and C5) dominate among species (Fig. 5d), suggesting an ancestral 8-chromosome karyotype of Brassicodae and Camelinodae (tAKI) where the analysis was largely sampled from. Additionally, the karyotype of *Thlaspi arvense* (basial lineage Aethionemoideae) were profiled as seven chromosomes, with only limited rearrangements relative to the tAKI. This was consistent with previous reports ^27^, demonstrating reliability of our *GouMang*.

**Fig. 5.**
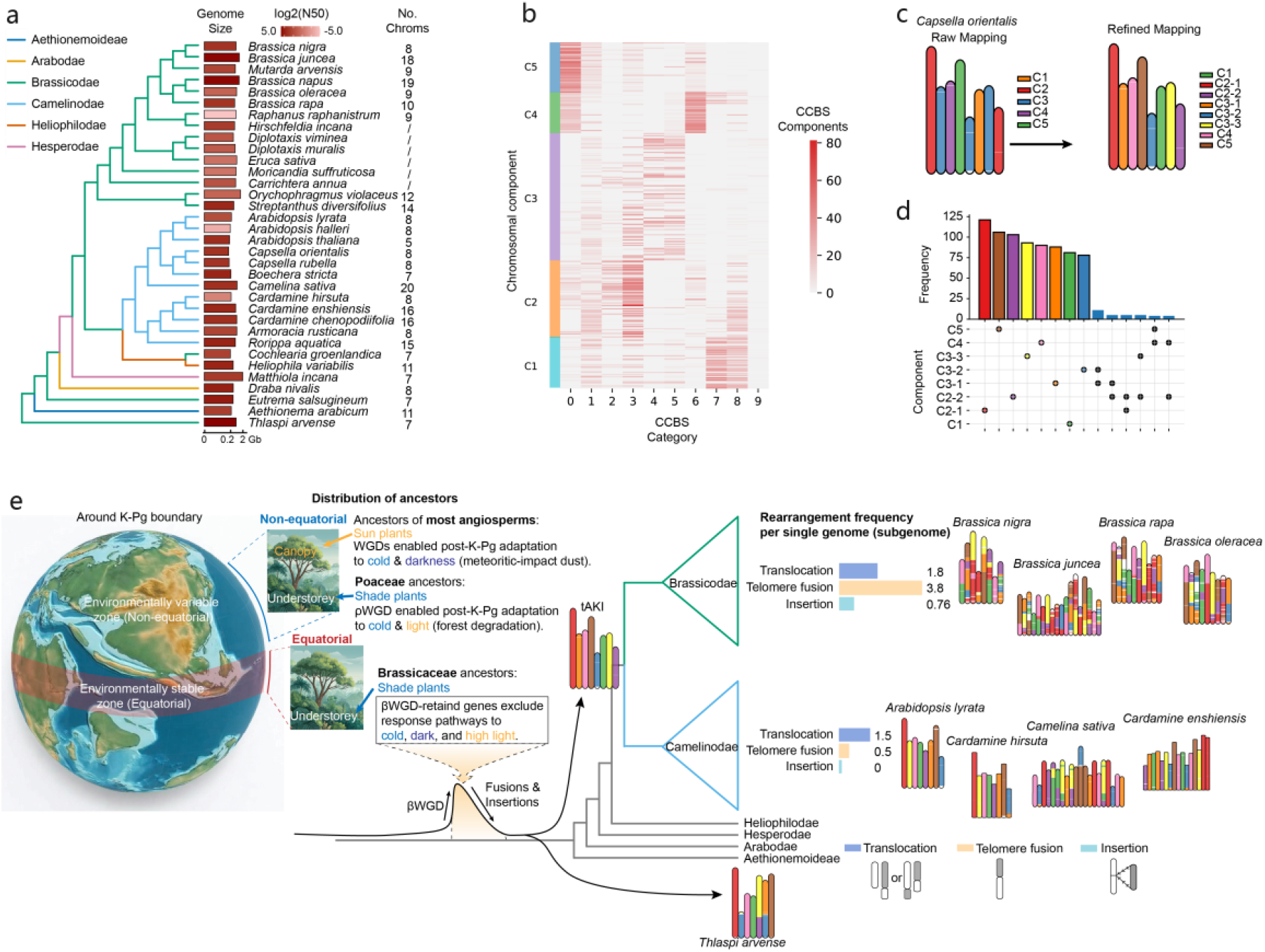
Karyotype evolution in Brassicaceae. **a**, Phylogenetic relationships of Brassicaceae species included in this study, with genome sizes and chromosome numbers indicated for each. **b**, Five common chromosomal components were grouped from ten CCBSs. **c**, The ancestral 8-chromosome karyotype of the two Brassicoideae supertribes (tAKI). **d**, Counts of chromosome with single or rearranged components across Brassicaceae species, revealing eight relatively conserved proto-chromosomes. **e**, Putative distribution of plant ancestors in ancient earth with karyotype diversification history in Brassicaceae.

The reconstruction of ancestral karyotype, further enabled to delineate the karyotypic dynamics within Brassicaceae (Fig. 5e). Brassicodae and Camelinodae exhibit distinct karyotype evolutionary modes: Camelinodae displays greater karyotypic stability, characterized by moderate translocation frequencies (~1.5 per genome/subgenome), rare telomere fusions (~0.5), and minimal chromosome insertions (~0), as exemplified by *Arabidopsis lyrata, Cardamine hirsuta, Camelina sativa*, and *Cardamine enshiensis*; In contrast, Brassicodae has undergone multiple whole-genome duplications (WGDs) and triplications (WGTs) ^51–53^ accompanied by more extensive chromosomal restructuring, with especially frequent telomere fusions (~3.8 per genome/subgenome) in species such as *Brassica nigra, B. juncea, B. rapa*, and *B. oleracea*. These findings highlight markedly different levels of karyotype diversification among lineages of Brassicaceae. Notably, comparing to grasses which rely predominantly on chromosome insertions, Brassicaceae frequently employ telomere fusions as chromosome number reduction mechanisms (Fig. 5e).

Additionally, intra-genome synteny analysis of *Capsella orientalis*, which retains a karyotype similar to the ancestral state, identified 18 proto-segment pairs generated by Brassicaceae-shared βWGD (Extended Data Fig. 11). Comparing to 12 common chromosomes in grasses fused from nine ρWGD-driven pairs, the βWGD may be followed by more extensive rearrangement (18 pairs to 8 common chromosomes). Notably, genes retained from the βWGD (Supplementary Table 5) showed no significant enrichment for cold- or dark-adaptation pathways commonly retained in WGDs which other angiosperm families shared ^41^, nor for the light-adaptation pathways enriched in the grass-shared ρWGD (Fig. 4e-g). This may reflect a divergent biogeographic, adaptative history of Brassicaceae compared to grasses and many other lineages (see Discussion).

## DISCUSSION

### *GouMang* Delineates Macroevolutionary Genic Karyotype Patterns across Plants

This study introduces *GouMang*, a novel approach for reconstructing comprehensive genic karyotype evolution across wide species through mining conserved chromosomal building blocks. These findings reveal a 12-chromosome karyotype of most recent grass ancestor, comprising the nine common chromosomal compositions as well as three high rearranged components. This ancestral karyotype subsequently diversified through lineage-specific chromosome reduce (via fusion or insertion) and increase (via WGDs), and itself originated from the ρWDG and subsequent rearrangements of nine earlier proto-chromosomes (EC1 to EC9). Parallel profiling in Brassicaceae identified an eight-chromosome tAKI, which diversified across major subfamilies within ranging levels following the shared βWGD. Notably, we observed distinct restructuring preferences between these lineages: Brassicaceae favored chromosome reduction via telomere fusions, whereas grasses more frequently utilized chromosomal insertions. These patterns underscore both the prevalence and the mechanistic diversity of karyotype dynamics in plant evolution.

Given that recent experimental work in *Arabidopsis* ^54^ has suggested that chromosomal rearrangements may promote reproductive isolation and accelerate speciation, the differential rates and modes of karyotype restructuring observed among lineages could reflect variations in evolutionary tempo or historical survival strategies. This opens several important questions for future research: Do lineages with extensive chromosomal restructuring evolve faster, potentially as a response to historical environmental shifts? While recently radiating groups such as Lake Victoria cichlids have provided evidence for the role of chromosomal rearrangements in explosive diversification ^13,14^. the contribution of ancient complex rearrangements to lineage divergence remains largely unexamined in highly rearranged plant taxa, particularly compared to well-studied, karyotypically stable groups such as primates ^55^. *GouMang* offers a scalable framework to address these questions in highly rearranged lineages, timely for the era of large-scale genome initiatives.

### WDGs Shaped Adaptive and Biogeographic History in Angiosperm lineages

Lineage-defining WGDs have been hypothesized to contribute to ancestral adaptation^41^. For instance, WGDs occurring around the K-Pg transition may have aided plants in coping with global cooling and meteorite impact driven darkness by retaining duplicated genes involved in cold and dark adaptation. Our analyses reveal contrasting ancestral stresses for early grasses and Brassicaceae, which appear to differ from those inferred for many other angiosperm lineages.

In grasses, genes retained after the ρWGD are enriched for pathways involved in cold and light stimuli responses, suggesting adaptation to cooler climates and more open, sun-exposed habitats. This signature is consistent with a post-K-Pg shift in non-equatorial regions, where forest decline created open niches that favored the transition of ancestral grasses from shade-tolerant understory plants to light-demanding sun plants (Fig. 5e). This adaptive shift may represent a pivotal event in the rise of modern grasses. Enhanced photomorphogenesis and the associated increase in biomass could have supported the expansion of herbivore ecosystems ^50^ and later the development of agricultural societies ^49^. We further propose that genes retained from the ρWGD likely served as a key genetic foundation for this shift (Fig. 4g).

In contrast, the Brassicaceae-shared βWGD shows no marked retention of genes linked to cold, dark, or light adaptation (Fig. 5). This divergent pattern implies a distinct biogeographic and adaptive history. The absence of cold-adaptation genes suggests that the ancestral Brassicaceae lineage likely occupied a relatively stable, equatorial region, a scenario aligned with “museum hypothesis” that the tropical regions served as stable refugia protecting from cold waves ^56–58^. Furthermore, the lack of light/dark adaptation genes suggests no major ecological transition between shade and sun niches during this period. Together, these lines of evidence point toward an ancestral equatorial understory niche for early Brassicaceae (Fig. 5e), a view that accords with phylogenetic reconstructions placing the family’s early evolution in tropical forest interiors before its crown-group radiated during the Eocene-Oligocene cooling-drying transition ^59,60^. The subsequent ecological success of Brassicaceae is often attributed to its “escape” from stable rainforests to exploit new niches opened by this global climatic shift ^59,60^.

Collectively, our findings illustrate a common macroevolutionary dynamic of plants, with WGDs periodically supplying a reservoir of adaptive genetic material and karyotype reshuffling potentially accelerating lineage divergence, explaining the distinct biogeographic histories of ancestor lineages.

### Leveraging Paleogenomic Information for Crop Improvement

Beyond revealing adaptive history, paleogenomic information may provide potential value in guiding future crop-improvement strategies. The set of genes retained from the grass ρWGD includes numerous regulators of key agricultural traits, many of which show functional and expression divergence between paralogous pairs (Fig. 4h). This genetic inventory offers a basis for systematically examining how ancestral WGDs may have contributed to modern crop domestication. Furthermore, *GouMang* could help infer biological roles for unexplored paralogous copies within conserved gene pairs. For instance, while *GS5* is a well-characterized regulator of grain size in rice ^61^ and its ρWGD-derived paralog remains functionally unstudied, prioritizing validation of such under-explored paralogs could accelerate identifications of contributors to yield and quality.

### Methodological Validation and Advantages of *GouMang*

*GouMang* yields more comprehensive reconstructions of whole evolutionary trajectory, comparing to previous approaches ^12,21,27^. The post-ρWGD karyotype in grasses (Extended Data Fig. 9a) and post-βWGD karyotype in Brassicaceae reconstructed by GouMang show high similarity to earlier models ^12,21,27^, supporting the methodological reliability. For the pre-ρWGD karyotype in grasses, however, our results differ from previously proposed five-to seven-chromosome models ^12,21^ (Extended Data Fig. 9b): some *GouMang* chromosomal components correspond to fused segments in prior reconstructions, while others appear split across multiple previously-proposed earlier proto-chromosomes. Re-examination of the three species (rice, sorghum, and *Brachypodium distachyon*) used in the seven-chromosome reconstruction (Extended Data Figs. 9c-d) indicates that these discrepancies arise from lineage-specific insertions in *B. distachyon*, which reflect recent fusions rather than the ancestral grass karyotype (Fig. 3k). By integrating much broader phylogenetic sampling, *GouMang* provides a more robust and comprehensive representation of the entire karyotype evolutionary trajectory.

The CCBS step of *GouMang* approach filters out evolutionarily “noisy” rearrangements while preserves core ancestral signatures, proving especially effective in deeply diverged, rearrangement-rich taxa. The organizational order of CCBS blocks along chromosomes shows remarkable variability across species in the same clusters, whereas CCBS compositions remain largely conserved (Fig. 2b). This pattern highlights the complexity of lineage-specific genomic reshuffling and supports our strategy of focusing on conserved chromosomal components, rather than attempting to resolve unique block orders for ancestral inference. It works by clustering chromosomes based on CCBS compositions rather than attempting to reconstruct precise block orders that are likely not unique.

## Acknowledgements

We thank Dr. Jianquan Liu (from Lanzhou University) and Dr. Pengchuan Sun (from Sichuan University) for the reviewing comments in Our *GouMang* framework and thank Dr. Hongtao Liu (from Shenzhen University) and Dr. Yuannian Jiao (Institute of Botany, Chinese Academy of Sciences) for the evaluation in the conclusions regarding grass adaptation to cooling and high-light conditions. We acknowledge the grants from National Natural Science Foundation of China [32350002, 32370257] and the Third Xinjiang Scientific Expedition Program [2022xjkk0200].

## Author contributions

J.G. and X.L. designed the *GouMang* algorithm. J.G. performed analyses. J.G., W.C., D.L., H.T., and X.L. interpreted data. J.G., H.T., and X.L. wrote the manuscript. X.L. supervised the study.

## Materials & Correspondence

This study did not generate new unique reagents.

